# The claustrum is required for reward acquisition under high cognitive demand

**DOI:** 10.1101/390443

**Authors:** Michael G. White, Chaoqi Mu, Hongkui Zeng, Brian N. Mathur

## Abstract

The claustrum is proposed to mediate a variety of functions ranging from sensory binding to top-down cognitive control of action, but direct functional assessments of this telencephalic nucleus are lacking. Here we employ the guanine nucleotide-binding subunit beta-4 cre driver line in mice to selectively monitor and manipulate claustrum projection neurons. Using fiber photometry, we find elevated claustrum activity prior to an expected cue during correct performance on a cognitively demanding five-choice response assay relative to a less-demanding one-choice version of the task. Claustrum activity during reward acquisition is also enhanced when cognitive demand is higher. Furthermore, we use optogenetic inhibition of claustrum prior to the expected cue to demonstrate that claustrum is critical for accurate performance on the five-choice, but not the one-choice, task. These results suggest the claustrum supports a cognitive control function necessary for reward acquisition under cognitively demanding conditions.

## Introduction

Across species, the claustrum is widely connected with the neocortex including sensory, motor, association and executive cortices (Crick and Koch 2005; Mathur 2014). This connectivity motivates a number of functional hypotheses (Remedios et al. 2010; Smith and Alloway 2010; Smythies et al. 2012; Mathur 2014; Patru and Reser 2015), including that the claustrum binds sensory information to generate conscious percepts (Crick and Koch 2005). Direct analysis of claustrum function is historically intractable, which necessitated indirect functional assessments of this structure. For instance, a recent study leveraged the dense anterior cingulate cortex input to claustrum (Smith and Alloway 2010; Wang et al. 2017; White et al. 2017) to show that activity of this circuit rises with, and is required for, optimal performance on a five-choice response task (White et al. 2018).

Novel transgenic tools, such as cre recombinase driver lines, now provide genetic access to the claustrum (Wang et al. 2017). In this study, we use the guanine nucleotide binding protein beta 4 cre driver line (GNB4-cre) for monitoring and manipulation of claustrum projection neurons in awake, freely moving mice. Given the role of the anterior cingulate cortex input to the claustrum on five-choice response task performance, we herein examine the role of the claustrum itself on this task. Understanding claustrum function stands to inform higher order brain functions involving functional coordination across the cortical mantle (Miller and Buschman 2013; Koch et al. 2016; Tononi et al. 2016; White and Mathur 2018a).

## Results

### Claustrum contributes to complex, but not simple, task performance

To confirm that claustrum projection neurons express cre recombinase in the GNB4-cre mouse, we injected a virus expressing eYFP in a cre-dependent manner into the claustrum of GNB4-cre mice (Figure 1A). We used parvalbumin (PV) immunostaining to delineate claustrum borders (Figure 1B) (Mathur et al. 2009; White et al. 2018) and observed that virus expression and PV immunostaining were isomorphic (Figure 1C). We next examined the morphological and electrophysiological identity of GNB4-positive (+) neurons by performing whole-cell recordings from labeled neurons using recording pipettes filled with AlexaFluor®-594. We found that GNB4+ neurons were spiny (Figure 1D), consistent with a projection neuron identity (Braak and Braak 1982; Hur and Zaborszky 2005; Watakabe et al. 2014). Basic membrane properties of these neurons are shown in Figure 1E and representative responses to current injection steps are shown in Figure 1F. Because capacitance delineates two claustrum projection neurons (White and Mathur 2018b), this membrane property can further identify GNB4+ neurons. In particular, we find a wide range of GNB4+ capacitance values (89 to 176 pF) consistent with sampling from both subtypes (type I = 118 ± 16 pF; type II = 158 ± 9 pF [mean ± SD]; White and Mathur 2018b).

**Figure 1:**
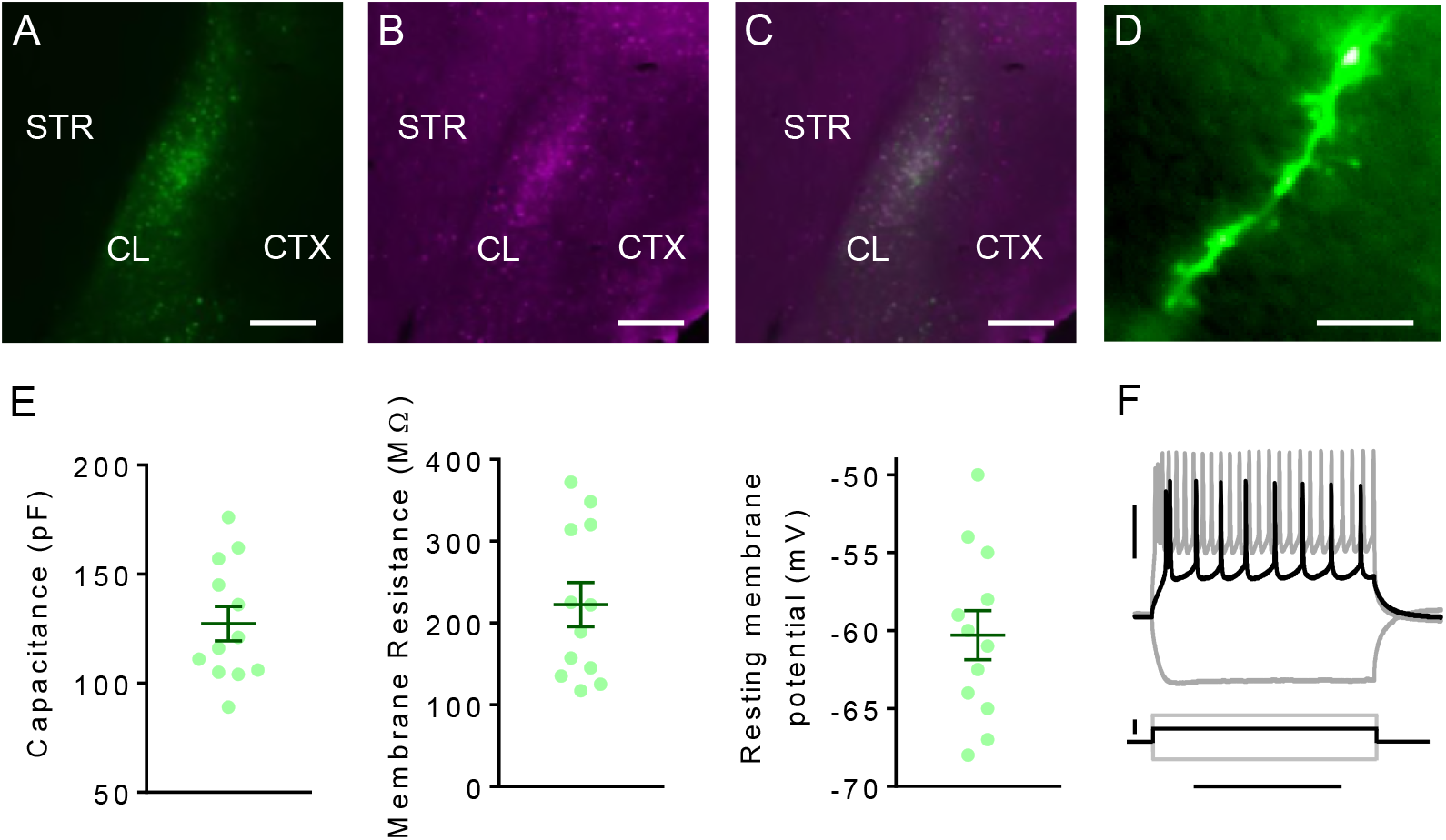
The GNB4-cre transgenic mouse line provides genetic access to claustrum projection neurons. (A) Cre-dependent viral expression of eYFP in mouse claustrum. (B) Immunostaining of claustrum parvalbumin (PV) expression. (C) Labeling of virally-expressed eYFP and immunolabeled PV were isomorphic. (D) Dendritic spines were evident upon whole-cell patch clamp records of GNB4-positive claustrum neurons using an internal solution including AlexFluor^®^-594. (E) Left: Mean capacitance from labeled neurons was 127 ± 9 (mean ± SD) pF. Middle: Mean membrane resistance was 222 ± 34 MΩ. Right: Mean resting membrane potential was −60 ± 2 mV. (F) Representative traces from labeled neuron in responsive to current injection steps. Horizontal scale bars = 200 μm (A-C), 10 μm (D), 400 ms (F). Vertical scale bars = 30 mV (F [top]), 200 pA (F [bottom]).

To determine if claustrum is critical for the five-choice serial reaction time task (5CSRTT) performance, we injected cre-dependent halorhodopsin (AAV-DIO-eNPhR3.0) or eYFP (AAV-DIO-eYFP) into the claustrum of GNB4-cre mice and implanted optical fibers to expose the claustrum to 470 nm light (Figure 2A). In acute brain slices, 470 nm light readily blocked action potential generation in claustrum neurons expressing eNPhR3.0 during a depolarizing voltage step (Figure 2B). Experimental and control mice were trained to perform the 5CSRTT and subsequently a one-choice stimulus-response task (1CSRTT; Figure 2C). The 1CSRTT was used to control for the basic sensory and motor components of the 5CSRTT. The claustrum was exposed to 470 nm light during the inter-trial interval (ITI) pseudo-randomly on 33% of 5CSRTT and 1CSRTT trials (Figure 2D). This protocol was derived from our previous work showing ACC input to claustrum is critical for optimal task performance within 1 s of cue onset (White et al. 2018). On 5CSRTT trials paired with 470 nm stimulation, AAV-DIO-eNPhR3.0 mice were less accurate compared to control trials; whereas no difference in accuracy was observed in AAV-DIO-eYFP mice (Figure 2E). Light delivery did not change the number of omissions or the response latency on correctly performed 5CSRTT trials in AAV-DIO-eNPhR3.0 or AAV-DIO-eYFP mice (Figure 2F and 2G). For AAV-DIO-eNPhR3.0 mice, we found that accuracy deficits on inactivation trials during the 1CSRTT were less than those on the 5CSRTT (Figure 2H). For AAV-DIO-eYFP mice, no differences in accuracy deficits were observed between the two tasks (Figure 2H).

**Figure 2:**
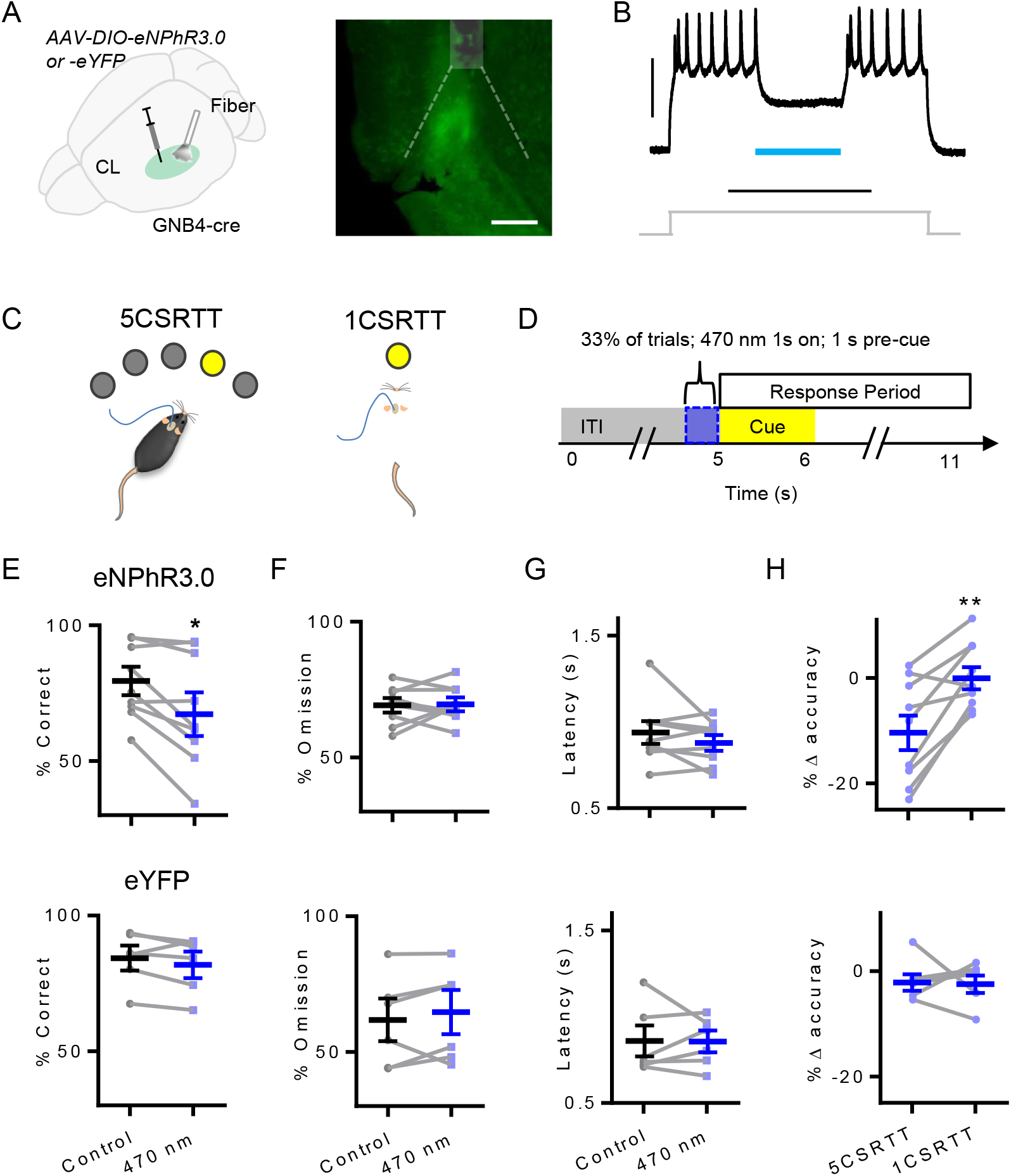
Inactivation of claustrum prior to cue presentation selectively disrupts response accuracy on the five-choice serial reaction time task (5CSRTT). (A) Left: Schematic illustrating that a virus expressing either halorhodopsin (n = 9) or eYFP (n = 6) in a *cre*-dependent manner (AAV-DIO-eNPhR3.0 or -eYFP, respectively) was injected bilaterally into the claustrum (CL) of GNB4-cre mice. Optical fibers were chronically implanted bilaterally in the claustrum. Right: Photomicrograph illustrating fiber optic implant (white box) position above claustrum, estimated light path (dotted lines), and viral expression. (B) Representative trace from a claustrum neuron expressing halorhodopsin (eNPhR3.0) that shows loss of action potential firing in the presence of 470 nm light during a depolarizing current injection. (C) Mice were trained to perform the 5CSRTT and subsequently a one-choice control task (1CSRTT). (D) Experimental schematic illustrating that 470 nm light was delivered to the claustrum during the inter-trial interval (ITI) 1 s prior to the onset of the cue on 33% of 5CSRTT or 1CSRTT trials. (E) Top: For eNPhR3.0 mice performing the 5CSRTT, choice accuracy was reduced on trials paired with 470 nm light delivery pre-cue compared to control trials. Paired *t* test, *t*(8) = 3.20, *P* = 0.013. Bottom: No changes in choice accuracy were found on 5CSRTT trials paired with light delivery in eYFP mice. Paired *t* test, *t*(5) = 1.38, *P* = 0.23. (F) Top: The omission rate was not different on 5CSRTT trials paired with 470 nm light delivery compared to control trials for eNPhR3.0 mice. Paired *t* test, *t*(8) = 0.14, *P* = 0.91. Bottom: Light delivery did not alter omission rate in eYFP mice. Paired *t* test, *t*(5) = 0.97, *P* = 0.38. (G) Top: The latency to correctly respond was not different on 5CSRTT trials paired with 470 nm light delivery compared to control trials for eNPhR3.0 mice. Paired *t* test, *t*(8) = 1.24, *P* = 0.25. Bottom: Light delivery did not alter latency to correctly respond in eYFP mice. Paired *t* test, *t*(5) = 0.042, *P* = 0.97. (H) Top: For eNPhR3.0 mice, the reduction in choice accuracy on the 5CSRTT resulting from 470 nm light delivery was significantly larger than the reduction during 1CSRTT performance. Paired *t* test, *t*(8) = 4.44, *P* = 0.0022. Bottom: For eYFP mice, there was no difference in the change in choice accuracy on the 5CSRTT resulting from 470 nm light delivery compared to the change during 1CSRTT performance. Paired *t* test, *t*(5) = 0.14, *P* = 0.89. Horizontal scale bars = 200 μm (A), 500 ms (B). Vertical scale bar = 30 mV.

To further exclude possible motor or reward-related effects elicited by claustrum inactivation, we performed a real-time place preference (RTPP) assay. We found that both AAV-DIO-eNPhR3.0 and AAV-DIO-eYFP mice did not exhibit a preference for either the control side of the chamber or the side of the chamber paired with 470 nm light stimulation (Figure 2 – Supplement 1A and 1B). Assessing whether claustrum inactivation affects movement, we found no difference in the velocity of AAV-DIO-eNPhR3.0 and AAV-DIO-eYFP mice before 470 nm light exposure compared to the 1 s of inactivation + the 1.5 s following inactivation (Figure 2 – Supplement 1C).

### Claustrum activity is sensitive to task load

We used a custom-made *in vivo* fiber photometry system to monitor calcium-dependent activity of claustrum in GBN4-cre mice injected with a virus expressing GCaMP6f in a cre-dependent manner (AAV-FLEX-GCaMP6f; Figure 3A). The system and data processing were designed to minimize sources of noise, such as fluctuations in excitation laser intensity, motion-related artifacts, and bleaching artifacts. Fluctuations in excitation laser intensity were controlled for by exciting a stable fluorophore, AlexaFluor^®^-488 (AF^®^-488; Figure 3 – Supplement 1A-1C). Motion-related artifacts were controlled for by multiplexing excitation of claustrum with 405 nm light, the isosbestic wavelength for GCaMP6 (Kim et al. 2016), and 473 nm light (Figure Supplement 1A and 1B). Regressing out the signals from 405 nm excitation and AF^®^-488 excitation attenuated noise (Figure 3 – Supplement 1D-1F). To control for photo-bleaching during 30 min 5CSRTT sessions, signals were examined in 10 trial bins.

**Figure 3:**
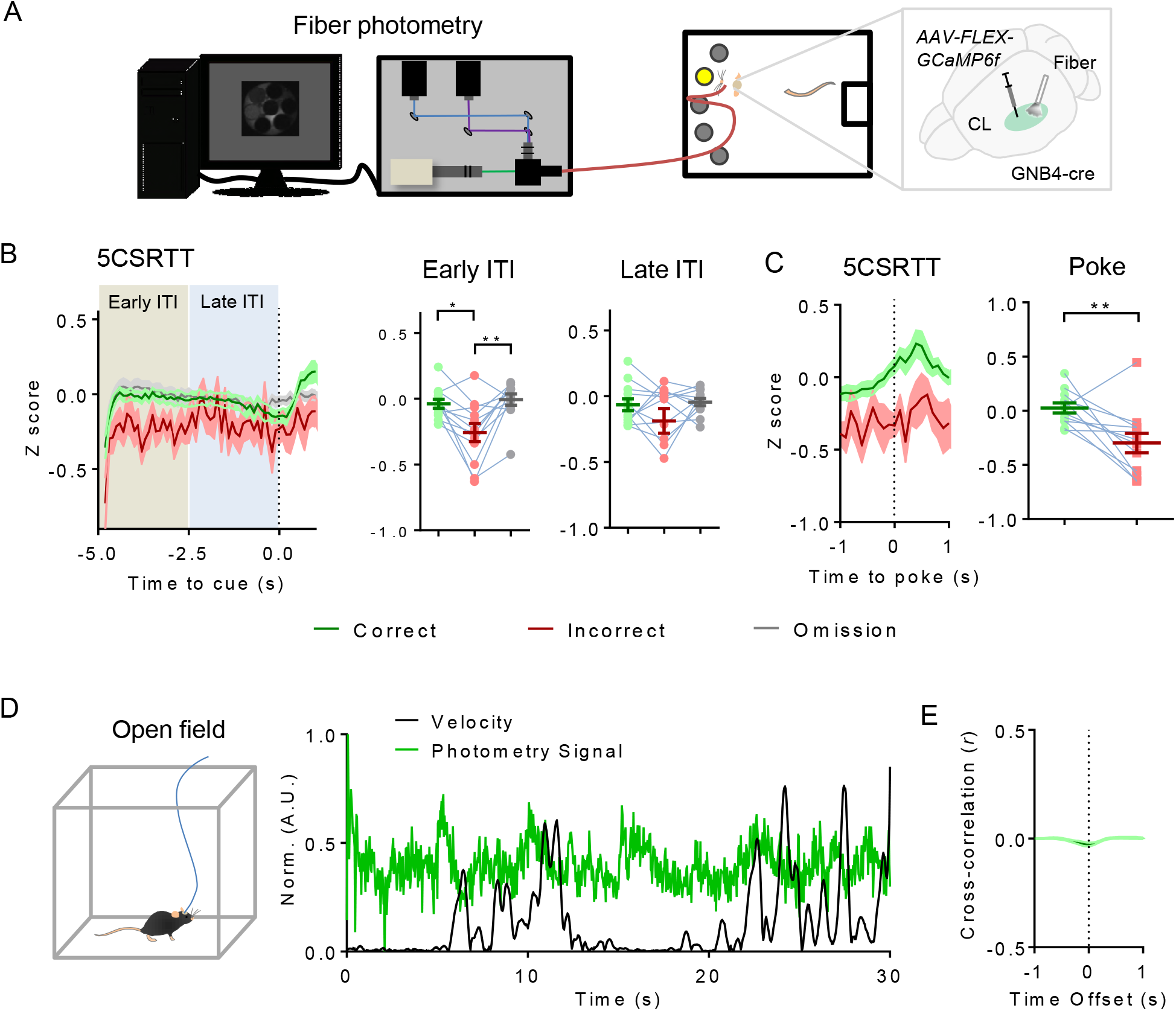
Claustrum activity during 5CSRTT performance and open field behavior. (A) Experimental schematic illustrating that claustrum activity was monitored during 5CSRTT performance in GNB4-cre mice injected with AAV-FLEX-GCaMP6f (n = 12). (B) Left: Average calcium-dependent activity of claustrum aligned to 5CSRTT cue onset is shown for correct (green), incorrect (red), and omission (gray) trials. Middle: Early in the ITI (5 to 2.5 s prior to cue onset), claustrum activity was greater for correct and omission trials relative to incorrect trials. Repeated measures ANOVA, *F*(2,11) = 9.48, *P* = 0.0014; post-hoc Tukey’s test, *P* = 0.015 (correct vs. incorrect), *P* = 0.0095 (omission vs. incorrect). Right: Late in the ITI (2.5 to 0 s prior to cue onset), claustrum activity did not differ across trial types. Repeated measures ANOVA, *F*(2,11) = 1.975, *P* = 0.17. (C) Left: Average claustrum activity aligned to correct or incorrect nose pokes during 5CSRTT performance is shown. Right: Average claustrum activity (1 s prior to 1 s after poke) was greater for correct nose pokes compared to incorrect nose pokes. Paired *t* test, *t*(11) = 3.14, *P* = 0.0095. (D) Left: Cartoon of mouse in an open field. Right: Representative traces of normalized claustrum activity (green) and velocity (black) during movement in an open field from GNB4-cre mice. (E) Cross-correlation between the photometry signal and movement velocity.

During 5CSRTT performance, claustrum activity was elevated on correct and omission trials in the early phase of the ITI relative to incorrect trials (Figure 3B). In the late phase of the ITI, there were no differences in claustrum activity among the different trial types (Figure 3B). We next aligned claustrum activity to correct and incorrect nose pokes during 5CSRTT performance. Average claustrum activity around the time of nose poke was significantly greater for correct nose pokes compared to incorrect nose pokes (Figure 3C). To assess if claustrum activity bears any relationship with movement, we monitored activity during free movement in an open field (Figure 3D). Claustrum activity was weakly and negatively correlated with movement velocity (Average *r* = −0.0062 Figure 3E). We next examined if claustrum activity during correct 5CSRTT performance reflects task load. To this end, we first compared claustrum activity during the ITI between correctly performed 5CSRTT and 1CSRTT trials (Figure 4A) and found relatively enhanced claustrum activity during the early ITI on the 5CSRTT but no differences during the late ITI (Figure 4B). We next compared the two tasks by aligning activity to the correct nose pokes (Figure 4C). We did not observe any activity differences between the two tasks immediately prior to or subsequent to nose pokes (Figure 4D). Lastly, we aligned claustrum activity to the acquisition of sucrose pellets (Figure 4E). Activity was greater for the 5CSRTT relative to the 1CSRTT immediately before and after acquisition of the sucrose pellet (Figure 4F).

**Figure 4:**
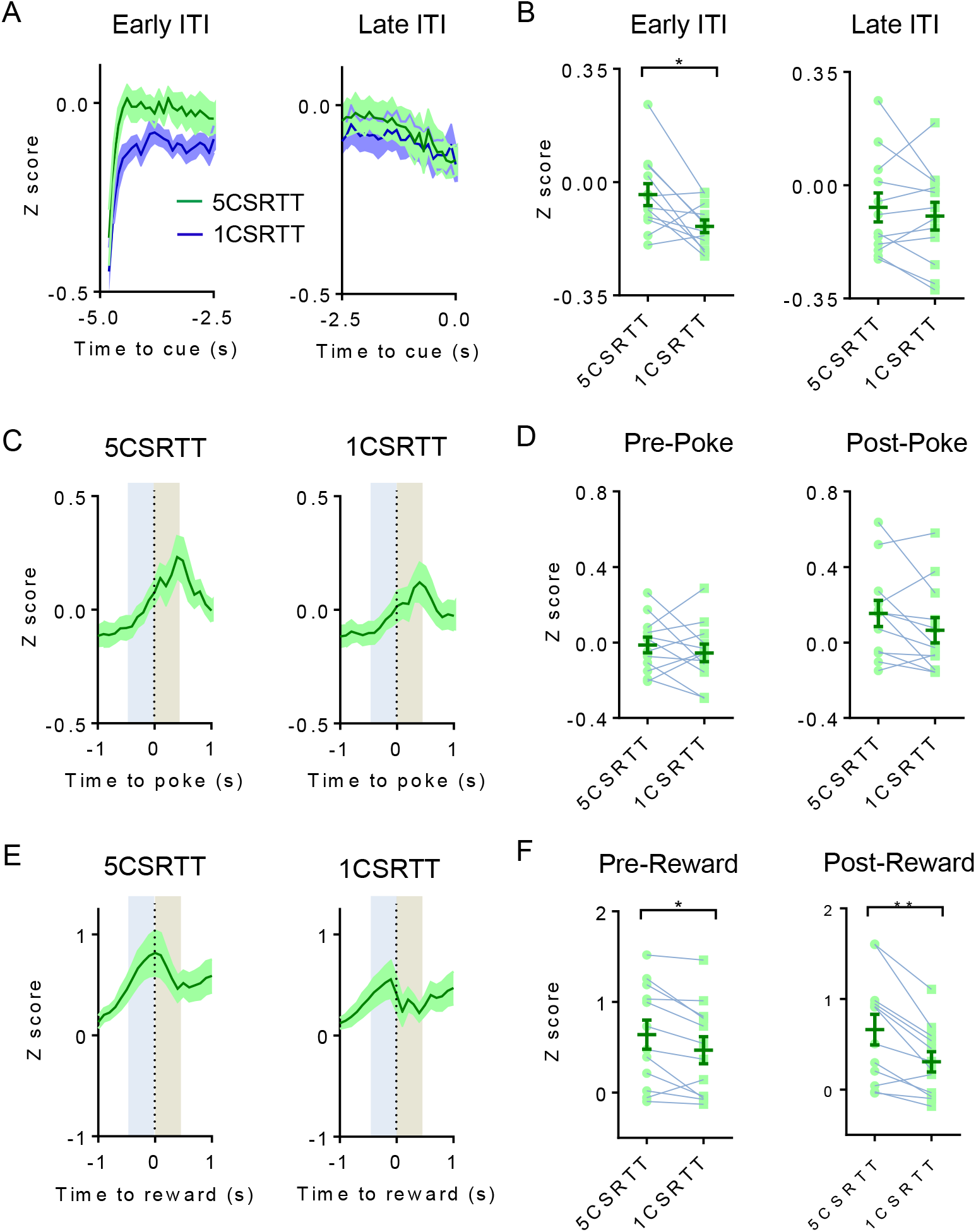
Claustrum activity on 5CSRTT versus 1CSRTT. (A) Left: Average claustrum activity during the early ITI for correct 5CSRTT trials (green) and correct 1CSRTT trials (blue). Right: Average activity during the late ITI for correct 5CSRTT and 1CSRTT trials. (B) Left: Claustrum activity during the early ITI was greater for correct 5CSRTT trials compared to 1CSRTT trials. Paired *t* test, *t*(11) = 2.92, *P* = 0.014. Right: no difference was observed between correct 5CSRTT and 1CSRTT trials during the late ITI. Paired *t* test, *t*(11) = 0.87, *P* = 0.41. (C) Left: Average claustrum activity aligned to correct nose pokes during 5CSRTT performance and right: during 1CSRTT performance. (D) Left: Claustrum activity prior to nose poke (average of 0.5 s prior, gray box) was not different between 5CSRTT and 1CSRTT. Paired *t* test, *t*(11) = 0.88, *P* = 0.40. Right: activity immediately subsequent to nose poke (average of 0.5 s subsequent, brown box) was not different between 5CSRTT and 1CSRTT. Paired *t* test, *t*(11) = 1.85, *P* = 0.09. (E) Left: Average claustrum activity aligned to collection of the reinforcement (sucrose pellet) on correct 5CSRTT trials and right: on correct 1CSRTT trials. (F) Left: Claustrum activity prior to collection of reward pellet (average of 0.5 s prior) was greater for the 5CSRTT relative to the 1CSRTT. Paired *t* test, *t*(11) = 3.11, *P* = 0.01. Right: activity immediately subsequent to collection (average of 0.5 s subsequent) was greater for the 5CSRTT compared to the 1CSRTT. Paired *t* test, *t*(11) = 4.41, *P* = 0.0010.

## Discussion

Our results suggest that claustrum is required for optimal performance on the cognitively-demanding 5CSRTT but not the 1CSRTT or open field movement. Claustrum activity on the 5CSRTT is higher on accurate compared to inaccurate task performance. In addition, relative to the 1CSRTT, claustrum activity on the 5CSRTT is higher at early stages prior to the cue and at reward acquisition. These findings are in line with the previous finding that the activity of anterior cingulate cortex inputs to claustrum are greater during 5CSRTT than 1CSRTT (White et al. 2018) and support the notion that the claustrum is involved in cognitive control of action.

It is important to note that our photometry analysis detects population level claustrum projection neuron activity. Therefore, it is difficult to determine, for example, whether increased activity-dependent calcium signals reflect recruitment of more claustrum neurons, an increase in firing of a subset of claustrum neurons, synchrony of claustrum neurons, or some combination of these possibilities. As such, future studies will need to assess if selective activation of functionally distinct subpopulations may explain our finding that claustrum activity on omission trials, as with correct trials, is elevated relative to incorrect trials during the 5CSRTT ITI, for example.

Claustrum activity may also be driven by retrosplenial cortex, which is heavily connected with claustrum (White et al. 2017), encodes disengagement from the external environment in humans (Buckner et al. 2008) and is part of a homologous task negative network in rodent (Lu et al. 2012). Given the claustrum’s widespread cortical interconnectivity and response to top-down command (White et al. 2018), the present results suggest a role for the claustrum in supporting cortical networks underlying cognitive control of goal-directed behavior under demanding conditions.

## Materials and methods

### Animals

34 GNB4-cre mice bred from a C57BL/6J background of both sexes were used (Wang et al. 2017). Mice used for electrophysiology were 10-20 weeks of age at the time of experiments and group-housed with food and water available *ad libitum*. In contrast, mice used for behavioral experiments were 16-30 weeks of age at the time of experiments. These mice were singly-housed, weighed daily, and fed daily to maintain 90% of *ad libitum* weigh. All mice were on a 12h light-dark cycle beginning at 0700 and 5CSRTT experiments were performed during the light cycle. For optogenetic experiments, control and experimental groups were comprised of mice from the same litters to minimize any litter effects. This study was performed in accordance with the National Institutes of Health Guide for Care and Use of Laboratory Animals and the University of Maryland, School of Medicine, Animal Care and Use Committee.

### Viral vectors and stereotaxic procedures

For optogenetic inhibition or *in vivo* fiber photometry monitoring of claustrum 80-110 nL of AAV vectors expressing *loxP-flanked* double inverted open reading frames (DIO) of halorhodopsin (AAV5-eF1a-DIO-eNPhR3.0-eYFP; University of Pennsylvania Vector Core) or GCaMP6f (AAV9-EF1a-DIO-GCaMP6f; University of Pennsylvania Vector Core), respectively, were injected bilaterally at two rostrocaudal levels of the claustrum (4 injections) in GNB4-cre mice. In control mice and mice used for whole-cell electrophysiology, the same approach was used but with a vector expressing eYFP (AAV5-eF1a-DIO-eYFP; University of Pennsylvania Vector Core). Relative to bregma, claustrum coordinates were 1) anterior-posterior: +1.34 mm, medial-lateral ± 2.3 mm, dorsal-ventral (from the brain surface): −2.35 mm; and 2) anterior-posterior: +0.86 mm, medial-lateral ± 2.75 mm, dorsal-ventral (from the brain surface): −2.55 mm. Viral incubation was no fewer than 3 weeks.

Mice used for *in vivo* optogenetics experiments were implanted bilaterally with chronic indwelling fiber optic implants into claustrum. Fiber optic implants were custom-made using high NA (0.66) fiber (Prizmatix Ltd; Giv’at Shmuel, Israel) epoxied into ceramic ferrules (ThorLabs Inc; Newton, NJ) and affixed to the skull with dental cement. Mice used for *in vivo* fiber photometry received a unilateral implant into claustrum, which was custom-made from low (0.22) NA fiber and ceramic ferrules (ThorLabs Inc; Newton, NJ). Fiber implant placement was confirmed post-hoc using immunohistochemistry. The light path angle (*θ*) was approximated to ensure accurate implantation using the known refractive index (*n*) of cortical tissue (Sun et al., 2012) and the respective fiber NA according to the following formula: NA = *n*sin*θ*. The distance of light penetration was approximated using previous estimates for each type of fiber (Cui et al., 2013, Al-Juboori et al., 2013).

### *Ex vivo* brain slice preparation for electrophysiology

Following anesthetization, mice were decapitated, the brains were extracted, and 250 μm coronal sections were sliced using a vibrating microtome in a high-sucrose artificial cerebrospinal fluid (aCSF). The aCSF was ice-cold, carbogen (95% O_2_, 5% CO_2_)-bubbled, and consisted of 194 mM sucrose, 30 mM NaCl, 4.5 mM KCl, 1 mM MgCl2, 26 mM NaHCO3, 1.2 mM NaH2PO4, and 10 mM D-glucose. Sections were incubated after slicing for 30 min at 33°C in carbogen-bubbled aCSF (315-320 mOsm) that contained 124 mM NaCl, 4.5 mM KCl, 2 mM CaCl_2_, 1 mM MgCl_2_, 26 mM NaHCO_3_, 1.2 mM NaH_2_PO_4_, and 10 mM D-glucose. Sections were incubated at room temperature until use for whole-cell patch-clamp recordings, and recordings were performed in the same aCSF formulation used for incubation.

### Whole-cell current and voltage-clamp recordings

Whole-cell recordings were performed at 29°C −31°C using borosilicate glass recording pipettes of 3-7 MΩ resistance. Recording pipettes were filled with a potassium-based solution (290-295 mOsm; pH 7.3) composed of 126 mM potassium gluconate, 4 mM KCl, 10 mM HEPES, 4 mM ATP-Mg, 0.3 mM GTP-Na and 10 mM phosphocreatine. Clampex software (version 10.4; Molecular Devices; Sunnyvale, CA) was used for electrophysiological recordings, which were filtered at 2 kHz and digitized at 10 kHz. Internal pipette solutions also contained hydrazide dye conjugated with AlexaFluor®-488 (40 μM) for visualization of dendritic spines. GNB4+ neurons expressing eYFP or halorhodopsin were identified using epifluorescence and targeted for recordings. For neurons expressing halorhodopsin, 470 nm light was delivered with an external LED for 300 ms during a 1 s current injection that elicited neuron firing to determine successful inhibition of firing using 470 nm light.

### Five-choice serial reaction-time task (5CSRTT)

Mice were trained to perform the 5CSRTT (Muir et al. 1996; Passetti et al. 2002; Dalley et al. 2004; Robinson et al. 2008) in operant chambers (Med Associates; St. Albans, VT) housed within sound-attenuating cabinets as previously described (White et al. 2018). Briefly, mice are trained to nose poke into one of five pseudo-randomly illuminated apertures (cue). Trials are preceded by a 5 s inter-trial interval (ITI) and responses are allowed during the cue and up to 5 s after the cue. Correct nose pokes resulted in a sucrose pellet dispensed into a receptacle on the wall opposite to the five apertures. A new trial ITI did not begin until 5 s after the reinforcement was collected. Nose pokes into the incorrect aperture, no response (omission), and nose pokes during the ITI resulted in a 5 s time out period, during which the house light was extinguished. Any nose poke during the time out period restarted the 5 s time out.

For optogenetic experiments, 470 nm light was delivered bilaterally during experimental sessions using an LED system (Plexon Inc; Dallas, TX); light delivery occurred pseudo-randomly on 33% of trials during the ITI period continuously for 1 s prior to the onset of the cue. A session ended after 100 trials or 30 min, whichever occurred first, and data was averaged across five sessions.

Mice were habituated, shaped, and trained before experiments as previously described (White et al., 2018). Mice were habituated to the operant chamber for two 15 min sessions with sucrose pellets available in the receptacle. Shaping occurred in two phases. First, all apertures were illuminated and any nose poke was reinforced. Apertures were initially loaded with sucrose pellets to facilitate initial poking. In the second phase, only one aperture was illuminated and only a nose poke into the illuminated aperture was reinforced. Mice progressed through each shaping phase after meeting criterion, which was defined as 30 correct nose pokes in a 30 min session. No time outs were given during shaping.

Mice were then progressed through two 5CSRTT training phases that were identical to the final 5CSRTT except for the cue duration. For the first training phase, the cue duration was 10 s and for the second training phase, the cue duration was 5 s. For training phases, criterion was defined as 60% accuracy and 30 responses within a session. Mice were over-trained on the final task (1 s cue) until choice accuracy reached a stable baseline before manipulations. Subsequent to 5CSRTT experimental sessions, mice were trained to perform a one-choice modification of the 5CSRTT (1CSRTT). All aspects of the task remained the same, except the middle aperture was illuminated and active on every trial. Mice were over-trained until performance stabilized before beginning experimental sessions. In 1CSRTT experimental sessions, 470 nm light was delivered as described above for the 5CSRTT continuously for 1 s prior to the onset of the cue.

### Real-time place preference (RTPP) and open field assay

The RTPP apparatus consisted of a two-sided chamber connected with a narrow corridor. The RTPP assay consisted of habituation and test sessions. In the habituation session, mice were placed initially in the narrow corridor of the chamber and allowed free exploration for 20 min. On the following day, mice performed the test session, which paired one side of the chamber with 1 s of continuous 470 nm light bouts. These bouts were repeated at most every 20 s in order to approximate the amount of stimulation that occurred during the 5CSRTT sessions. The amount of time spent in each side of the chamber during sessions and ambulatory velocity was recorded with EthoVision XT v 11.5 (Noldus, Wageningen, The Netherlands). The velocity during the 0.5 s of light delivery and the subsequent 2 s was compared to the 2.5 s preceding light delivery to determine any effect of light delivery on movement. The same apparatus was used as an open field to measure calcium-dependent activity of claustrum during movement using *in vivo* fiber photometry.

### *In vivo* fiber photometry

Photometry data from 5CSRTT and open field experiments were collected using a customized *in vivo* fiber photometry system. Two single wavelength laser modules were used (Opto Engine LLC; Midvale, Utah), a 473 nm laser for optimal GCamP6f excitation, and a 405 nm laser to excite GCaMP6f at its isosbestic wavelength (Kim et al. 2016). As such, emission from 405 nm excitation was used to control for image artifacts due to photometry cable motion, background fluorescence, and other sources of noise (Kim et al. 2016). The two lasers were multiplexed at 10 Hz, resulting in a continuous 20 Hz pulse train. Both laser beams were directed using the two-bounce method into a dichroic filter cube optimized for 473 and 405 nm excitation, as well as for 510 nm emission (Chroma Technology Corporation; Bellow Falls, VT). The two excitation wavelengths were focused through a 4X fluorite objective (Olympus; Tokyo, Japan) onto a multimode fiber bundle that consisted of seven individual multimode fibers. One fiber was connected to the mouse via chronic multimode fiber implant, while another fiber was placed inside a tube of AlexaFluor^®^-488 (40 μM) to control for variability in laser energy. Emissions from GCaMP6f and AlexaFluor^®^-488 through the multimode fibers were detected as an image of the fiber bundle using the ORCA Flash 4.0LT high-resolution CMOS camera (Hamamatsu Photonics KK; Hamamatsu City, Japan). Laser multiplexing and image acquisition were synchronized using an Arduino Leonardo microcontroller and time-locked to 5CSRTT trials when necessary. Camera image acquisition parameters were controlled through the HCImage Software for Hamamatsu cameras.

### Data analysis and statistics

5CSRTT photometry data were analyzed using a combination of MATLAB (Mathworks; Natick, Massachusetts) and Prism v 6.0.1 (GraphPad Software; La Jolla, CA). MATLAB Graphical User Interface (GUI) codes were written specifically to process the photometry raw image datasets, each one consisting of up to 40,000 images. First, the raw signal from the photometry signal fiber and the fluorophore control fiber was extracted by selecting ROIs for both fibers, and then the pixel values were averaged across the entire ROI to extract the raw signal from the image. The raw signals were then subtracted from a corresponding background signal from the same image, which is the baseline transmission of ambient light from a photometry fiber when excitation lasers are off. The processed signals were then sorted based on if the image was taken during a 473 or 405 nm laser pulse. Further processing required a separate MATLAB GUI for linear regression and averaging each individual dataset across all trials. First, 473 and 405 nm photometry signals were regressed with the corresponding control fluorophore signal, and the residuals of the regression were then used for further processing. The 473 nm signal was then regressed with the 405 nm signal as a covariate, and the residuals of the regression were extracted as the fully processed photometry signal from the claustrum. The processed signal was then sorted based on trial type (correct, incorrect or omission), and averaged across trials for each specific trial type, and then across all five 5CRSTT runs for each animal. Further statistical analysis was performed in GraphPAD Prism. All statistics are displayed as mean ± standard error unless otherwise noted.

---

**Figure 2 – Supplement 1:**
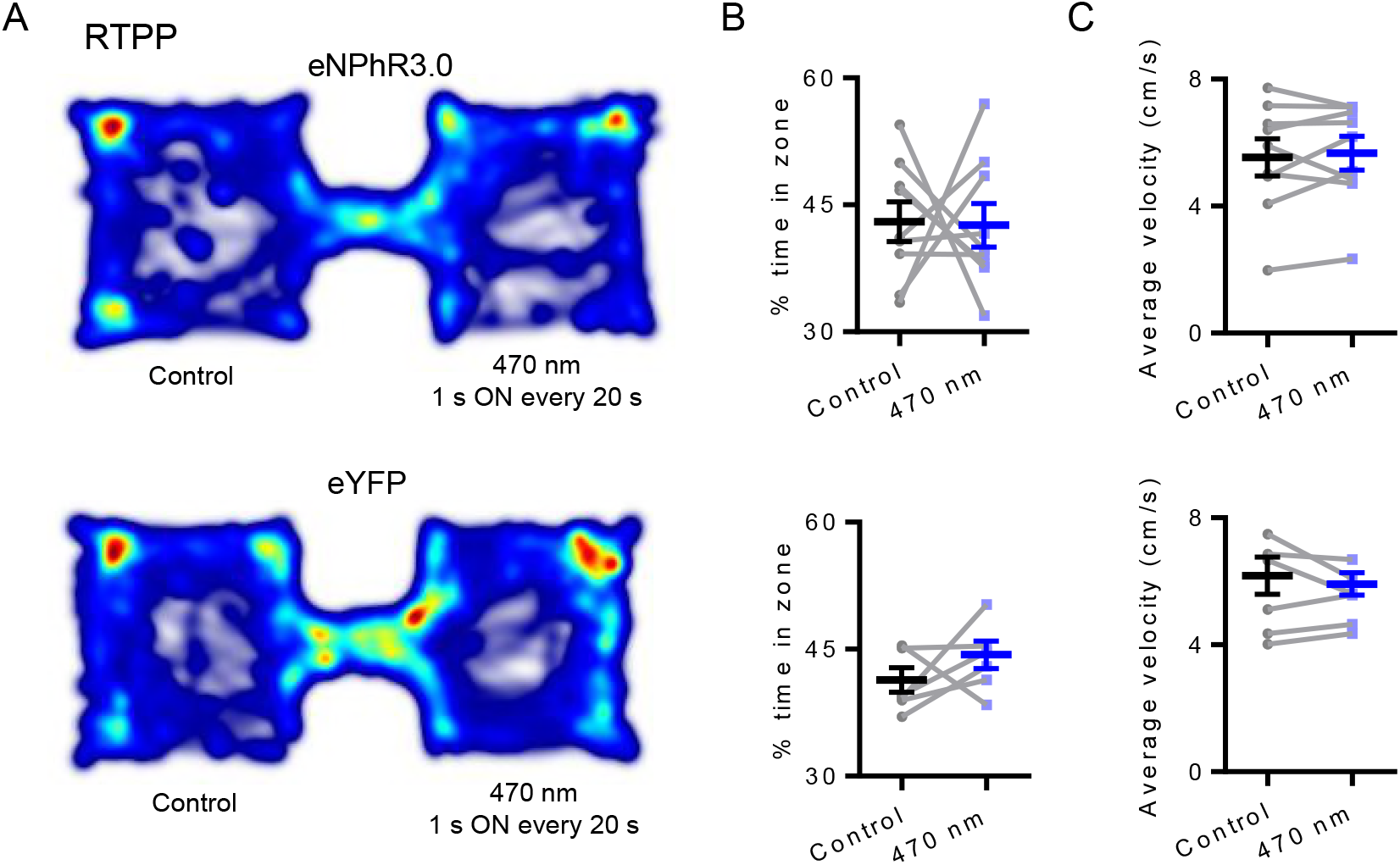
Claustrum inactivation is not reinforcing and does not affect movement. (A) Top: Representative heat map from a GNB4-cre mouse expressing eNPhR3.0 in claustrum during a real-time place preference assay (RTPP). One side of the chamber was paired with 1 s 470 nm light delivery to claustrum that cycled at a maximum rate of once every 20 s and the other side was not paired with light. Hot colors indicate more time spent in a given location and cool colors indicate less time spent in a given location. Bottom: Representative heat map from a GNB4-cre mouse expressing eYFP in claustrum during the RTPP assay. (B) Top: For eNPhR3.0 mice, no difference was observed in the time spent on the side paired with 470 nm light compared to the control side. Paired *t* test, *t*(8) = 0.089, *P* = 0.93. Bottom: For eYFP mice, no difference was observed in the time spent on the side paired with 470 nm light delivery to claustrum compared to the control side. Paired *t* test, *t*(5) = 1.20, *P* = 0.28. (C) Top: Movement velocity during the 2.5 s interval prior to 470 nm light delivery was compared to velocity during the 1 s of light delivery plus the 1.5 s interval following the offset of light delivery. For eNPhR3.0 mice, no differences in velocity were observed between the two intervals. Paired *t* test, *t*(8) = 0.55, *P* = 0.60. Bottom: For eYFP mice, no differences in velocity were observed between the two intervals. Paired *t* test, *t*(5) = 0.81, *P* = 0.45.

**Figure 3 – Supplement 1:**
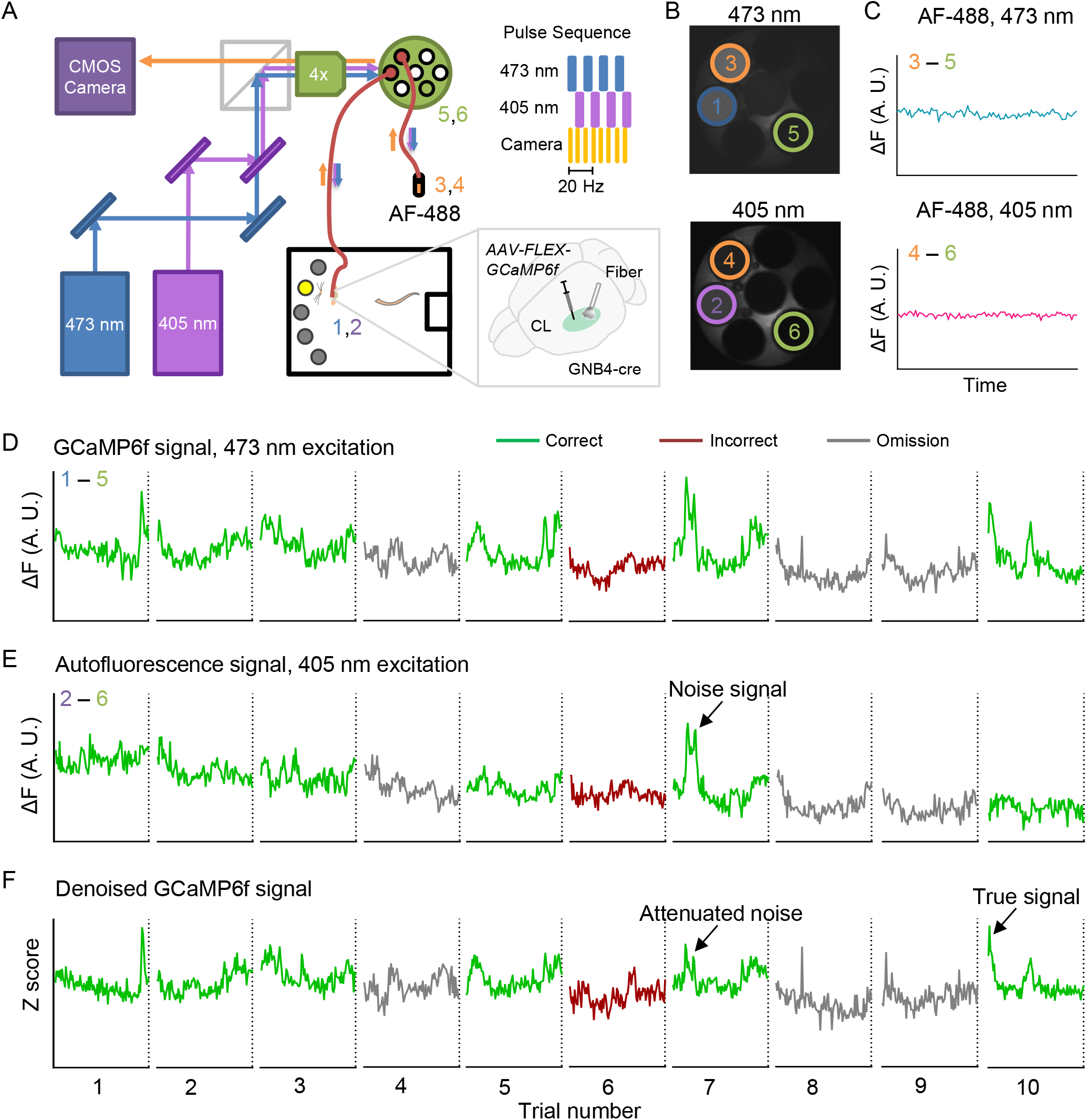
Custom *in vivo* fiber photometry system and data processing. (A) Schematic illustrating the custom *in vivo* fiber photometry system used for monitoring claustrum activity. Two excitation lasers, 473 nm (odd numbered signals) and 405 nm (even numbered signals), were multiplexed at 10 Hz. The beams were aligned, directed into a dichroic filter cube, and passed through a 4X objective into a multi-mode fiber bundle. One fiber was coupled to a chronic fiber implanted in the claustrum (signals numbered 1 and 2) of GNB4-cre mice injected with a cre-dependent virus expressing GCaMP6f (AAV-FLEX-GCaMP6f; inset). A control fiber was placed inside a tube of AlexaFluor^®^-488 (AF-488) fluorophore (signals numbered 3 and 4). GCaMP6f and AF-488 emissions collected by the fiber bundle were imaged with a Hamamatsu CMOS camera. The camera captured images at 20 Hz synchronized to each laser pulse. A fiber uncoupled to a fluorescence source (signals numbered 5 and 6) was used to measure background signals and obtain ΔF values. (B) Top: Representative image showing the fiber bundle in response to 473 nm excitation. Bottom: Representative image showing the fiber bundle in response to 405 nm excitation. (C) Top: Representative trace illustrating AF-488 ΔF values in response to 473 nm excitation (3 – 5). Bottom: Representative trace illustrating AF-488 ΔF values in response to 405 nm excitation (4 – 6). (D) Representative traces illustrating GCaMP6f ΔF values measured from claustrum with 473 nm excitation (1 – 5) from a bin of 10 5CSRTT trials that include different trial types. (E) Representative traces illustrating autofluorescence ΔF values measured from claustrum with 405 nm excitation (2 – 6). Arrow indicates example of a noise signal. (F) Representative traces illustrating denoised GCaMP6f signals obtained following regression of the AF-488 signal (C, top) and autofluorescence signal (E). Arrows indicate an attenuated noise signal and a true signal.

## Acknowledgments

This work was supported by National Institute on Alcohol Abuse and Alcoholism grants K22AA021414 and R01AA024845 (B.N.M.), Whitehall Foundation grant 2014-12-68 (B.N.M.), National Institute of General Medical Sciences grant T32GM008181 (M.G.W.), National Institute of Neurological Disorders and Stroke grant T32NS063391 (M.G.W.), and National Institute of Mental Health grant F31MH112350 (M.G.W.). The authors also are grateful for the assistance of Dr. Christof Koch and the Allen Institute for Brain Science.

